# Characterization of Pacific oyster (*Crassostrea gigas*) proteomic response to natural environmental differences

**DOI:** 10.1101/460204

**Authors:** Yaamini R. Venkataraman, Emma Timmins-Schiffman, Micah J. Horwith, Alexander T. Lowe, Brook Nunn, Brent Vadopalas, Laura H. Spencer, Steven B. Roberts

## Abstract

Global climate change is rapidly altering coastal marine ecosystems important for food production. A comprehensive understanding of how organisms will respond to these complex environmental changes can come only from observing and studying species within their natural environment. To this end, the effects of environmental drivers — pH, dissolved oxygen content, salinity, and temperature — on Pacific oyster (*Crassostrea gigas*) physiology were evaluated in an outplant experiment. Sibling juvenile oysters were outplanted to eelgrass and unvegetated habitat at five different estuarine sites within the Acidification Nearshore Monitoring Network in Washington State, USA to evaluate how regional environmental drivers influence molecular physiology. Within each site, we also determined if eelgrass presence that buffered pH conditions changed the oysters’ expressed proteome. A novel, two-step, gel-free proteomic approach was used to identify differences in protein abundance in *C. gigas* ctenidia tissue after a 29 day outplant by 1) identifying proteins in a data independent acquisition survey step and 2) comparing relative quantities of targeted environmental response proteins using selected reaction monitoring. While there was no difference in protein abundance detected between habitats or among sites within Puget Sound, *C. gigas* outplanted at Willapa Bay had significantly higher abundances of antioxidant enzymes and molecular chaperones. Environmental factors at Willapa Bay, such as higher average temperature, may have driven this protein abundance pattern. These findings generate a suite of new hypotheses for lab and field experiments to compare the effects of regional conditions on physiological responses of marine invertebrates.

## Introduction

Global climate change will influence estuarine dynamics and impact the organisms that inhabit these environments. Estuaries are already variable across spatial and temporal scales in terms of phytoplankton production (Pennock & Sharp 1986), nutrient availability (Paerl et al. 2014), heavy metal contamination (Liu et al. 2015), salinity (Banas et al. 2004), and carbonate chemistry (Feely et al. 2010; Pelletier et al. 2018). Since climate change will affect these parameters, it is important to consider how estuarine organisms will respond.

Proteomics, or the study of protein abundance and expression, can be used to shed light on physiological changes on a molecular level. Proteins direct all major cellular functions, thus examining protein abundance provides direct evidence of an organism’s physiological response to the estuarine environment (Tomanek 2014). The proteome is dynamic, as it must rapidly respond to perturbation, providing mechanistic information that standard gene expression and mRNA quantification studies cannot (Veldhoen et al. 2012; Flores-Nunes et al. 2015). As a result of the proteome’s dynamic nature, proteins analyzed at the time of collection represent an organism’s response to the environment in near real-time. Long-term exposure to environmental conditions, as well as natural organismal aging, are also reflected in the proteome (Hercus et al. 2003). Discovery-based proteomic methods can elucidate responses to environmental drivers (Flores-Nunes et al. 2015). Several studies have connected protein abundances with changes in laboratory-simulated environmental conditions, identifying key proteins and mechanisms involved in specific environmental responses (Timmins-Schiffman et al. 2014; Dineshram et al. 2016; Meng et al. 2017). While these studies provide insight into organismal adaptation and physiology, laboratory studies alone cannot fully encapsulate the effects of multiple environmental drivers within an ecosystem context (Riebesell & Gattuso 2014).

Although challenging, *in situ* field studies provide a necessary biological realism when considering variable environments (Slattery et al. 2012; Cornwall & Hurd 2016). Such experiments can be leveraged to study the effects of multiple environmental drivers on organismal physiology and to incorporate realistic variability, as opposed to examining the effect of a single stressor on an organism (Riebesell & Gattuso 2014). Through transcriptomics, Chapman et al. (2011) demonstrated the power of an *in situ* experimental design for examining the impacts of regional environmental conditions on Eastern oyster *(Crassostrea virginica)* physiology. Transcript signatures from *C. virginica* sampled from various locations in southeastern United States revealed that temperature, pH, salinity, dissolved oxygen and pollutant load at each location impacted gene expression. Furthermore, they were able to disentangle the interactions of these environmental factors on gene expression. RNA and protein abundances can be influenced by several environmental factors, and *in situ* studies can determine which drivers will be more important to consider for organismal physiology.

Marine invertebrates have proven to be informative bioindicators in proteomic studies to examine the effects of *in situ* conditions on organismal physiological responses to environmental change. When marine invertebrates have been exposed to varying environmental conditions, proteomics have demonstrated changes in cellular defense, immune responses, and genome function (Veldhoen et al. 2012). Changes in protein abundance in bivalves like the Pacific oyster (*Crassostrea gigas*) and blue mussels (*Mytilus edulis spp*.) have been used to develop biomarkers for environmental contaminants (Slattery et al. 2012; Beyer et al. 2017). Proteomic responses to natural environmental drivers have also been evaluated in bivalves. For example, shotgun proteomic analysis of *M. edulis* ctenidia from Baltic Sea microcosms revealed that low salinity conditions lead to decreased abundance of cytoskeleton proteins, as well as calcium-binding messenger calmodulin, which plays an important role in signalling and intracellular membrane trafficking pathways (Campos et al. 2016). Using a growing wealth of genomic information to understand how these species fare under differential environmental conditions is critical for monitoring natural populations and commercial aquaculture.

Pacific oyster (*Crassostrea gigas*) rearing in estuarine environments in Washington State, USA (WA) provides an ideal system to examine the effect of *in situ* environmental conditions on the expressed proteome. *C. gigas* are extensively farmed in two different estuarine systems that show substantial regional variation: Puget Sound and Willapa Bay. Puget Sound is a complex estuarine system with interconnected sub-basins, each with different freshwater inputs, residence times, and stratification levels (Feely et al. 2010; Bianucci et al. 2018; Pelletier et al. 2018). Willapa Bay is a large shallow estuary on the Pacific coast that exchanges approximately half its water volume with the Pacific Ocean at each tide (Banas et al. 2004, 2007). Seasonality and location within Puget Sound dictates temperature, dissolved oxygen, salinity, and pH conditions, while Willapa Bay conditions are influenced by diurnal fluctuations and proximity to either the ocean or rivers draining into the bay (Banas et al. 2007; Feely et al. 2010; Ruesink et al. 2015).

Both Puget Sound and Willapa Bay also host eelgrass beds (*Zostera spp*.) that affect environmental conditions, such as oxygen concentrations, on diurnal time scales. The 2012 Washington State Blue Ribbon Panel on Ocean Acidification outlines key early actions, which include the examination of “vegetation-based systems of remediation” (Action 6.1.1) to improve local pH through photosynthetic drawdown of carbon dioxide. This experiment set out to test whether protein abundance patterns reflect reduced stress within vegetation. For example, eelgrass beds may reduce emersion stress relative to unvegetated areas through shading, the retention of water, and increased evaporative cooling at low tide. They can also ameliorate effects of ocean acidification through photosynthetic activity. Reduced pathogen prevalence has also been documented in seagrass beds, but not specifically in eelgrass (Lamb et al. 2017). In contrast, eelgrass beds may drive more extreme carbonate chemistry conditions (Pacella et al. 2018). Lowe et al. (2018) also found that *C. gigas* shell strength and survival was significantly lower in eelgrass habitats in WA. Understanding how different aquaculture grow-out locations and habitats will affect the oyster’s ability to persist through environmental change is crucial for the industry and the ecosystem.

The purpose of this study was to use proteomic techniques to uncover the impacts of environmental drivers on Pacific oyster physiological outcomes in estuarine environments in WA. Naturally existing environmental variation was harnessed by outplanting *C. gigas* in different locations within Puget Sound and Willapa Bay, and habitat effects were taken into consideration by placing oysters in eelgrass and unvegetated habitats. Gel-free proteomic methods were used to examine the effects of outplant conditions on relative quantities of all expressed proteins in a series of *in situ* experiments in order to identify differentially abundant proteins. We predicted that differences in environmental drivers at each outplant location and within outplant habitats would yield unique protein abundance patterns. Oysters at outplant locations with warmer water temperatures, more variable water temperatures, lower dissolved oxygen content, lower salinity, or lower pH may have higher abundances of proteins related to environmental response. Eelgrass beds were expected to ameliorate stressful conditions, resulting in lower abundances of environmental stress response proteins than oysters in unvegetated habitats.

## Methods

### Shellfish Deployment

Sibling juvenile *C. gigas* (average shell length = 27.2 mm, age = 2 months) were outplanted for 29 days starting June 19, 2016 at five locations: Case Inlet (CI), Fidalgo Bay (FB), Port Gamble Bay (PG), Skokomish River Delta (SK), and Willapa Bay (WB) in Washington State, USA (Table 1). These sites were selected for differences in environmental parameters, as well as for the presence of unvegetated areas and eelgrass beds within each site. All sites were part of the Acidification Nearshore Monitoring Network, a network of sensors placed in various WA locations to monitor marine chemistry (ANeMoNe; Washington Department of Natural Resources). Prior to the outplant, oysters were reared in a controlled hatchery setting. At each site and habitat combination, custom-built Durafet-based sensors (Honeywell) were used to monitor pH. Commercially-available MiniDOT loggers (Precision Measurement Engineering) were used to measure dissolved oxygen, and Odyssey loggers were used to measure conductivity. All sensors recorded temperature measurements, and all sensors logged at 10-minute intervals across the outplant period, with the exception of SK, where sensors were installed two days into the outplant period. At each site, juvenile oysters were placed in bags of five oysters each directly onto the substrate at a tidal height of -1.5 MLLW, both inside and outside of eelgrass beds (n=15 per habitat type), for a total of thirty outplanted oysters per site. The animals were uniformly placed less than a lateral distance of 0.5 m from the sensors at the same tidal height as the instruments. Oysters were housed in exclusion cages to prevent predation. Juvenile oysters remained at each site for a 29-day exposure period. Because the ctenidia is the primary site where oysters interact with the environment, ctenidia samples were dissected at the end of the outplant and held on dry ice until storage at −80°C (Beyer et al. 2017; Meng et al. 2017).

**Table 1.** Latitude and longitude of *C. gigas* outplants, as well as time of day and tidal height at collection. Oysters were placed at five locations sites: Case Inlet (CI), Fidalgo Bay (FB), Port Gamble Bay (PG), Skokomish River Delta (SK), and Willapa Bay (WB).

| Location | Latitude | Longitude | Time at Collection | Tidal Height (MLLW, feet) at Collection |
| --- | --- | --- | --- | --- |
| CI | 47.3579367 | -122.7957627 | 12:15 | -1.8 |
| FB | 48.481691 | -122.58353 | 12:12 | -1.7 |
| PG | 47.842676 | -122.583832 | 11:11 | -1.6 |
| SK | 47.35523 | -123.1572 | 11:51 | -1.8 |
| WB | 46.4944789 | -124.0261356 | 9:28 | -1.7 |

Environmental data was treated as follows. Conductivity observations were removed when less than zero, which occurs when the instrument is dry at low tide. Remaining observations were converted to salinity measurements using the *swSCTp* function in the *oce* package in R (Kelley & Richards 2018; R Core Team 2018), with temperature at 25°C and pressure at 10 dbar. For dissolved oxygen, pH, and salinity datasets, data were removed when collected by probes 1) during low tide or 2) when tidal depth was less than one foot to remove readings where the probes may have been exposed. Values collected during low tide or a depths less than one foot were retained for temperature datasets. Outliers were screened using the Tukey method for temperature, dissolved oxygen, pH, and salinity datasets (Hoaglin et al. 1986). Uniform outplant tidal heights were checked using *prop.test* in R (R Core Team 2018).

A non-metric multidimensional scaling analysis (NMDS) was used to evaluate differences in environmental parameters. First, mean and variance were calculated for each day of the outplant. Values were log-transformed, and a separate Gower’s distance matrix was calculated for daily mean and daily variances, accounting for missing data. The NMDS was conducted with the Gower’s distance matrix to visually compare means or variances between sites and habitats. Significant differences between site and habitat were identified using a One-way Analysis of Similarities (ANOSIM) for each environmental parameter. Pairwise ANOSIM tests for significant one-way ANOSIM results and two-way ANOSIM tests by site and habitat were not conducted due to lack of replicates within each site-habitat combination. R Scripts are available in the supplementary Github repository (Venkataraman et al. 2018).

### Protein Discovery

To identify select protein targets for characterization across locations and environmental conditions, a subset of tissue samples were analyzed with data independent acquisition (DIA) mass spectrometry analysis. Two tissue samples were used from each site to make a peptide library and maximize the amount of protein abundance data collected from each sample.

#### Protein Quantification

Tissue samples were homogenized in a solution of 50 mM NH_4_HCO_3_ with 6M Urea (500μl). Tissues were then sonicated 3 times (Fisher Scientific Sonic Dismembrator Model 100) for 10 seconds each and cooled between sonications in a bath of ethanol and dry ice. Protein quantities were measured with the Pierce BCA Protein Assay Kit microplate assay with a limited quantity of sonicated sample (11 μL). The protein concentration was measured via spectroscopy at 540 nm in a Labsystems (Waltham, MA) Multiskan MCC/340 and accompanying Ascent Software Version 2.6. Protein concentrations were calculated based on a standard curve with BSA (Pierce) per manufacturer’s instructions.

#### Protein Digestion

Protein digestion followed the protocol outlined in Timmins-Schiffman et al. (2013). To each sample of 30 μg protein,1.5 M Tris pH 8.8 buffer (6.6 μL) and 200 mM TCEP (2.5 μL) were added. After solvent additions, each sample’s pH was verified to be basic (pH ≥ 8), and placed on a 37°C heating block for one hour. lodoacetamide (200 mM, 20 μL) was then added to each sample to maximize digestion enzyme access to protein cleavage sites. Samples were covered with aluminum foil to incubate in the dark for 1 hour at room temperature. Afterwards, diothiolthreitol (200 mM, 20 μL) was added and samples were incubated at room temperature for one hour. Lysyl Endopeptidase (Wako Chemicals) was then added to each sample in a 1 μg enzyme:30 μg oyster protein ratio, followed by one hour of incubation at room temperature. Urea was diluted with NH_4_HCO_3_ (25 mM, 800 μL) and HPLC grade methanol (200 μL). Trypsin (Promega) was added to each sample in a 1 μg trypsin: 30 μg oyster protein ratio for overnight digestion at room temperature.

#### Peptide Isolation

After overnight incubation, samples were evaporated to near dryness at 4°C with a speedvac (CentriVap ® Refrigerated Centrifugal Concentrator Model 7310021). Samples were then reconstituted in 100 μL of a 5% Acetonitrile and 0.1% Trifluoroacetic Acid (Solvent A) to isolate peptides. If samples were not at pH ≤ 2, 10-20 μL aliquots of 10% Formic Acid were added until this pH was achieved.

Before desalting peptide samples, Macrospin C18 columns (The Nest Group) were prepared by adding 200 μL of a 60% Acetonitrile with 0.1% Trifluoroacetic Acid (Solvent B). The columns were spun for three minutes at 2000 rpm, and flow-through liquid from the column was discarded. The spinning and discarding process was completed a total of four times. To wash columns, 200 μL of Solvent A was added to each column. The columns were once again spun for three minutes at 2000 rpm and liquid was discarded afterwards; the solvent addition, spinning, and discarding process was completed a total of three times.

To bind peptides to the columns, digested peptides were added to prepared columns, then the columns were spun at 3000 rpm for three minutes. The filtrate was pipetted back onto the column and spun again at 3000 rpm for three minutes. Solvent A (200 μL) was added to each column three separate times, then the column was spun for three minutes at 3000 rpm to wash salts off the column.

Peptides were eluted with two additions of 100 μL Solvent B to each column. Columns were spun at 3000 rpm for three minutes and the peptide fraction (filtrate) was reserved. Samples were placed in a speed vacuum at 4°C until they were nearly dry (approximately two hours) to dry peptides. Peptides were reconstituted with 60 μL of 3% Acetonitrile + 0.1% Formic Acid, and stored at −80°C.

#### Internal Standard Addition

Peptide Retention Time Calibration (PRTC; Pierce) is used as an internal standard to ensure consistency of peptides detected and measured throughout a mass spectrometry run. The stock solution of PRTC was diluted to 0.2 pmol/μL using 3% Acetonitrile with 0.1% Formic Acid. In a clean centrifuge tube, 6 μg of oyster protein and 0.376 pmol of PRTC were mixed together as per the PRTC user guide. Sample volume was brought up to 15 μL using a 3% acetonitrile and 0.1% formic acid solution. A quality control solution was also prepared (1 μL PRTC + BSA:3 μL 3% Acetonitrile and 0.1% Formic Acid solution).

### Data Independent Acquisition Mass Spectrometry

Peptides were analyzed on an Orbitrap Fusion Lumos mass spectrometer (Thermo Scientific) using Data Independent Acquisition Mass Spectrometry (DIA). DIA analyses were completed as a comprehensive, non-random analytical method for detecting peptide ions present within a sample to create a peptide library. The peptide library was then leveraged to develop a targeted proteomics assay for quantification (see Selected Reaction Monitoring Assay). A 30 cm analytical column and 3 cm pre-column were packed in-house with 3 μm C18 beads (Dr. Maisch). Samples were run in a randomized order. A blank injection followed each sample, with procedural blanks run at the very end. Every injection was 3 μL, which included 1 μg of oyster protein and 0.0752 pmol of PRTC. Peptides were analyzed in MS1 over the *m/z* range of 450-950 with 12 *m/z* wide windows with 5 *m/z* overlaps (Egertson et al. 2013). MS1 resolution was 60000 and AGC target was 400000 with a three second cycle time. The MS2 loop count was set to 20 and MS2 data was collected with a resolution of 15000 on charge state of 2 with an AGC target of 50000. No dynamic exclusion was used.

### Peptide-Centric Proteomic Analyses

Unknown peptide spectra from mass spectrometry samples were matched with known peptides using Peptide-Centric Analysis in the PECAN software (Ting et al. 2015). Raw mass spectrometry files were converted to mzML files, then demultiplexed using MSConvert (Chambers et al. 2012). The *C. gigas* proteome was digested with in silico tryptic digest using Protein Digestion Simulator (Riviere et al. 2015). All known peptides from the mzML files were identified in comparison to the digested *C. gigas* proteome (Riviere et al. 2015).

The PECAN-generated spectral library (.blib) file was used to detect peptides of interest in raw DIA files in Skyline (MacLean et al. 2010). Skyline identified peptides using chromatogram peak picking, where ions that elute at the same time and mass are detected as a peptide (file available at Panorama Public). All PRTC peptides and approximately 100 different oyster proteins and their peptide transitions were manually checked for retention time and peak area ratio consistency to determine a Skyline auto peak picker error rate (24.3% ± 25%, range: 0% to 100%).

Proteins had to satisfy four criteria to be considered appropriate targets for the study. 1) After an extensive literature search, functions related to oxidative stress, hypoxia, heat shock, immune resistance, shell formation, growth, and cellular maintenance were determined useful for evaluating environmental response. Proteins with annotations matching these functions were considered potential targets. 2) Protein data was then evaluated in Skyline to ensure there was no missing data for any peptide or sample. 3) Peaks with significant interference from other peptides were not considered. 4) Proteins needed at least two peptides with three transitions per peptide to quality as a potential target for downstream assays. The fifteen proteins (41 peptides and 123 transitions) that matched all of these criteria were selected as targets (Table 2).

**Table 2.** Proteins used as targets for a Selected Reaction Monitoring Assay (SRM). Targets were identified based on differential abundance and stress-related annotations. At least two peptides and six transitions were included in the assy for each protein. The protein Catalase had two isoforms under separate proteome IDs — CHOYP_CATA.1.3|m.11120 and CHOYP_CATA.3.3|m.21642 — and target peptides were chosen from both IDs. A total of four peptides and twelve associated transitions were used as SRM targets.

| Protein | Proteome ID | Number of Peptides | Number of Transitions |
| --- | --- | --- | --- |
| 3-ketoacyl-CoA<br>thiolase | CHOYP_ACAA2.<br>1.1\|m.30666 | 3 | 9 |
| Peroxiredoxin-5 | CHOYP_BRAFLD<br>RAFT_119799.1.<br>1\|m.23765 | 2 | 6 |
| Thioredoxin<br>reductase 3 | CHOYP_BRAFLD<br>RAFT_122807.1.<br>1\|m.3729 | 3 | 9 |
| Protein phosphatase<br>1B | CHOYP_BRAFLD<br>RAFT_275870.1.<br>1\|m.12895 | 3 | 9 |
| Carbonic anhydrase<br>2 | CHOYP_CAH2.1.<br>1\|m.42306 | 3 | 9 |
| Catalase 1 | CHOYP_CATA.1.<br>3\|m.11120 | 3 | 9 |
| Catalase 2 | CHOYP_CATA.3.<br>3\|m.21642 | 1 | 3 |
| Glucose-6-phosphate<br>1-dehydrogenase | CHOYP_G6PD.2.<br>2\|m.46923 | 3 | 9 |
| Heat shock 70 kDa<br>protein | CHOYP_HS12A.<br>25.33\|m.60352 | 2 | 6 |
| Heat shock protein 70<br>B2 | CHOYP_HSP74.<br>1.1\|m.13095 | 2 | 6 |
| NAD(P)<br>transhydrogenase | CHOYP_LOC100<br>633041.1.1\|m.354<br>28 | 2 | 6 |
| Glycogen<br>phosphorylase | CHOYP_LOC100<br>883864.1.1\|m.417<br>91 | 3 | 9 |
| Multidrug resistance-<br>associated protein | CHOYP_MRP1.5.<br>10\|m.34368 | 2 | 6 |
| Protein disulfide-isomerase 1 | CHOYP_PDIA1.1<br>.1\|m.5297 | 3 | 9 |
| Protein disulfide-isomerase 2 | CHOYP_PDIA3.1<br>.1\|m.60223 | 3 | 9 |
| Puromycin-sensitive<br>amidase | CHOYP_PSA.1.1\|<br>m.27259 | 3 | 9 |

### Selected Reaction Monitoring Assay

Following the protein discovery phase (DIA), proteins were isolated as described above from an additional five randomly selected samples per site and habitat combination (for a total of 5 oysters per group) and analyzed with Selected Reaction Monitoring (SRM). Samples were prepared as described for DIA, except tissue samples were homogenized in 100 μL, and peptide samples were evaporated at 25°C after peptide isolation.

Proteins of interest identified from the DIA analysis were used as targets in a SRM assay following the workflow and informatic pipeline of (Timmins-Schiffman et al. 2016). Target peptide transitions were monitored using SRM on a Thermo TSQ Vantage. SRM data were collected during a gradient of 2-60% acetonitrile over 40 minutes. All samples were run in technical duplicates in a randomized order with a 1 μg oyster peptide and 0.0752 pmol PRTC injection. A quality control injection and blank injection were run after every five sample injections, and PRTC peptides were monitored throughout the experiment.

#### Target Peptide Specificity

To ensure SRM assay specificity to oyster peptides of interest, oyster peptides were diluted in a background matrix of similar complexity (Pacific geoduck — *Panopea generosa* — peptides), then analyzed using the oyster SRM assay. An oyster-specific SRM target would decrease in abundance with a decreasing abundance of oyster peptides in a mixture. Non-specific peptides — more likely to be found in background matrix of similar complexity — or peptides susceptible to interference would not correlate with oyster peptide abundance, and therefore, would be uninformative. Five *C. gigas* samples used for SRM were randomly selected and pooled in equal quantities. A ten-sample oyster:geoduck dilution series was prepared and run using the same methods as other SRM samples.

#### Target Analysis

Raw SRM files, a background *C. gigas* proteome, and the PECAN spectral library file from DIA were used to create a Skyline document (file available at Panorama Public). Correct transition peaks were selected based on predicted retention times from DIA results by comparing the relative retention times between identical PRTC peptides in the DIA and SRM datasets (R^2^ = 0.99431). Based on peptide specificity analyses, heat shock protein 70 B2 and one constituent peptide of glucose-6-phosphate 1-dehydrogenase were removed from analyses (S11).

Further filters were applied to the data to maintain only high quality peptides and transitions in the analysis. Coefficients of variation were calculated between technical replicates for each peptide transition. Peptides were removed from the dataset if CV > 20%. To maintain high sample quality, any sample missing data for more than 50% of peptide transitions was deemed poor quality for downstream analyses and excluded. Abundance data was normalized using total ion current (TIC) values from the mass spectrometer. Consistency between technical replicates was verified in remaining samples using a NMDS with TIC-normalized data and a euclidean dissimilarity matrix. Technical replicates were consistent if replicates lay closer together than to other samples in the NMDS. These replicates were then averaged for multivariate analytical methods.

Averaged technical replicate data was used to determine if peptides were differentially abundant between outplant sites and habitats. Before proceeding with analysis, peptide abundances were subjected to a Hellinger transformation to give low weights to any peptides with low counts. A NMDS was used to visually compare relative peptide abundance. One-way ANOSIM tests by site, region (Puget Sound vs. Willapa Bay), and habitat, as well as a two-way ANOSIM test by site and habitat, were used to determine significant differences. Pairwise ANOSIM tests and post-hoc similarity percentage (SIMPER) analyses were conducted for each one- or two-way ANOSIM result significant at the 0.05 level. The first ten SIMPER entries were deemed influential peptides for each significant comparison.

The importance of environmental variables for explaining peptide abundance was evaluated with a redundancy analysis (RDA). For each site and habitat combination, mean and variance were calculated for pH, dissolved oxygen, salinity, and temperature over the course of the entire outplant. Environmental variables were then used as predictors to constrain peptide abundance. Predictors with missing values were not included. A triplot was used to visually assess differences in peptide abundance by site and habitat and the influence of individual peptides and environmental parameters. Analysis of Variance (ANOVA) was used to calculate significance of the RDA and environmental variables, with predictors deemed significant at p < 0.05. Since estuarine sites are highly variable, a second RDA was conducted constraining peptide abundance by environmental conditions on the day of collection to evaluate robustness of proteomic methods. R Scripts are available in the supplementary Github repository (Venkataraman et al. 2018).

## Results

### Conditions at Outplant Locations

Outplanted oysters experienced environmental variables representative of standard summer conditions in Puget Sound (PS) and WB. Wild or cultured oysters were present at the same tidal elevation as sensors, so outplant conditions represent experiences of non-experimental organisms. NMDS plots revealed mean environmental conditions were more similar among sites than variances. Both NMDS had stress values less than 0.20 and were considered appropriate multivariate representations of environmental data (Mean NMDS: stress = 0.0170, p = 0.0100; Variance NMDS: stress = 0.0340, p = 0.0100). Mean dissolved oxygen and temperature were significantly different between sites (Table 4; ANOSIM; dissolved oxygen: R = 0.4063, p = 0.0530; temperature: R = 1.0000, p = 0.0020), but not between habitats. Variances of all environmental parameters were significantly different between sites (Table 5; ANOSIM; pH: R = 0.5313, p = 0.0180; dissolved oxygen: R = 0.6800, p = 0.0030; salinity: R = 0.8400, p = 0.0130; temperature: R = 0.9400, p = 0.0010). WB had the warmest, but least variable, temperature of 18.0°C ± 1.3°C compared to averages ranging from 14.8°C ± 1.8°C to 16.1°C ± 1.7°C in PS (Table 3, Figure 2). All variance values, mean pH, and mean salinity were not significantly different between habitats (Table 4, Table 5).

**Figure 1.**
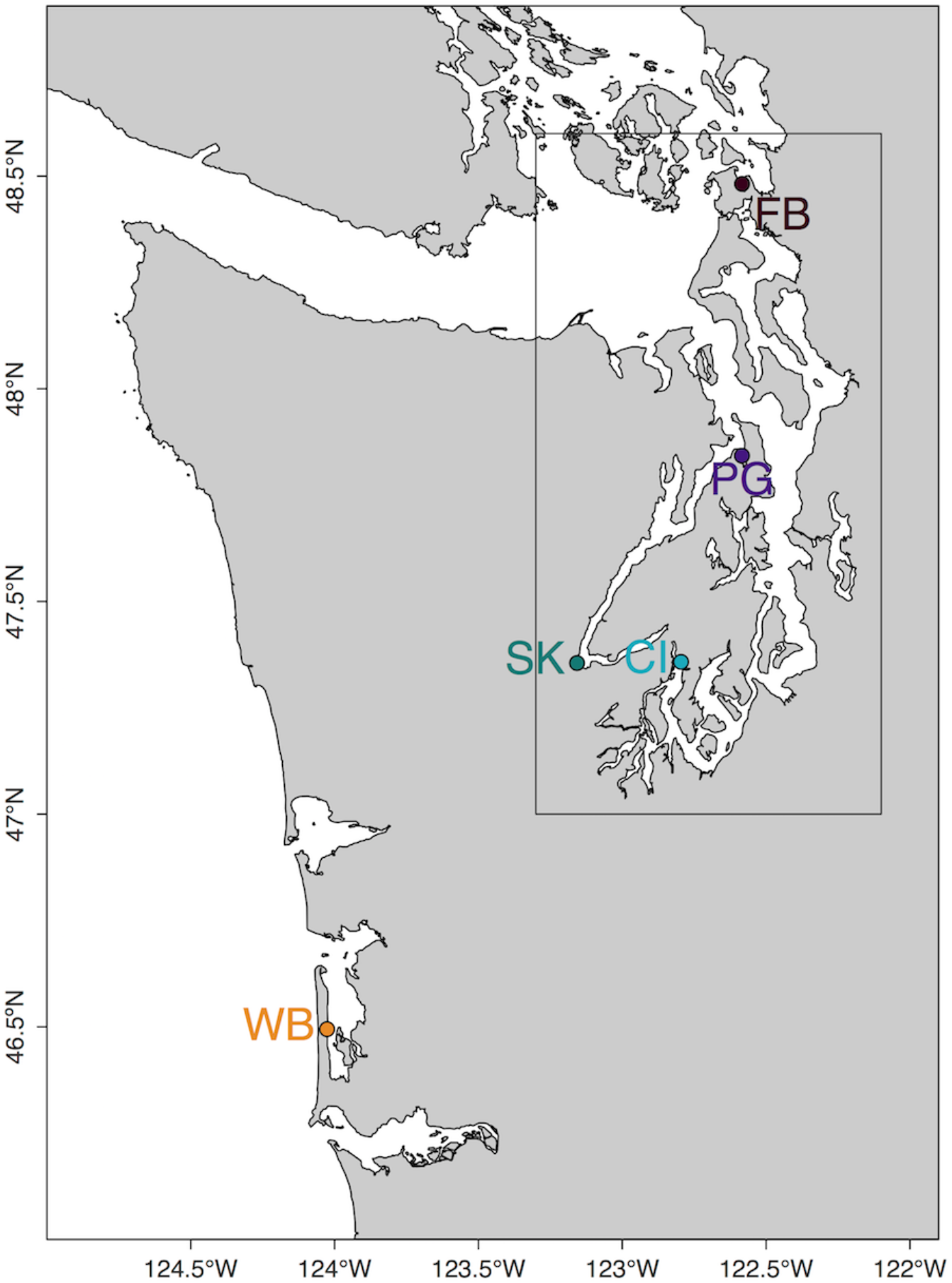
Map of outplant locations. Oysters were placed at five sites: Case Inlet (CI), Fidalgo Bay (FB), Port Gamble Bay (PG), Skokomish River Delta (SK), and Willapa Bay (WB). Sites in Puget Sound are outlined in a grey box.

**Figure 2.**
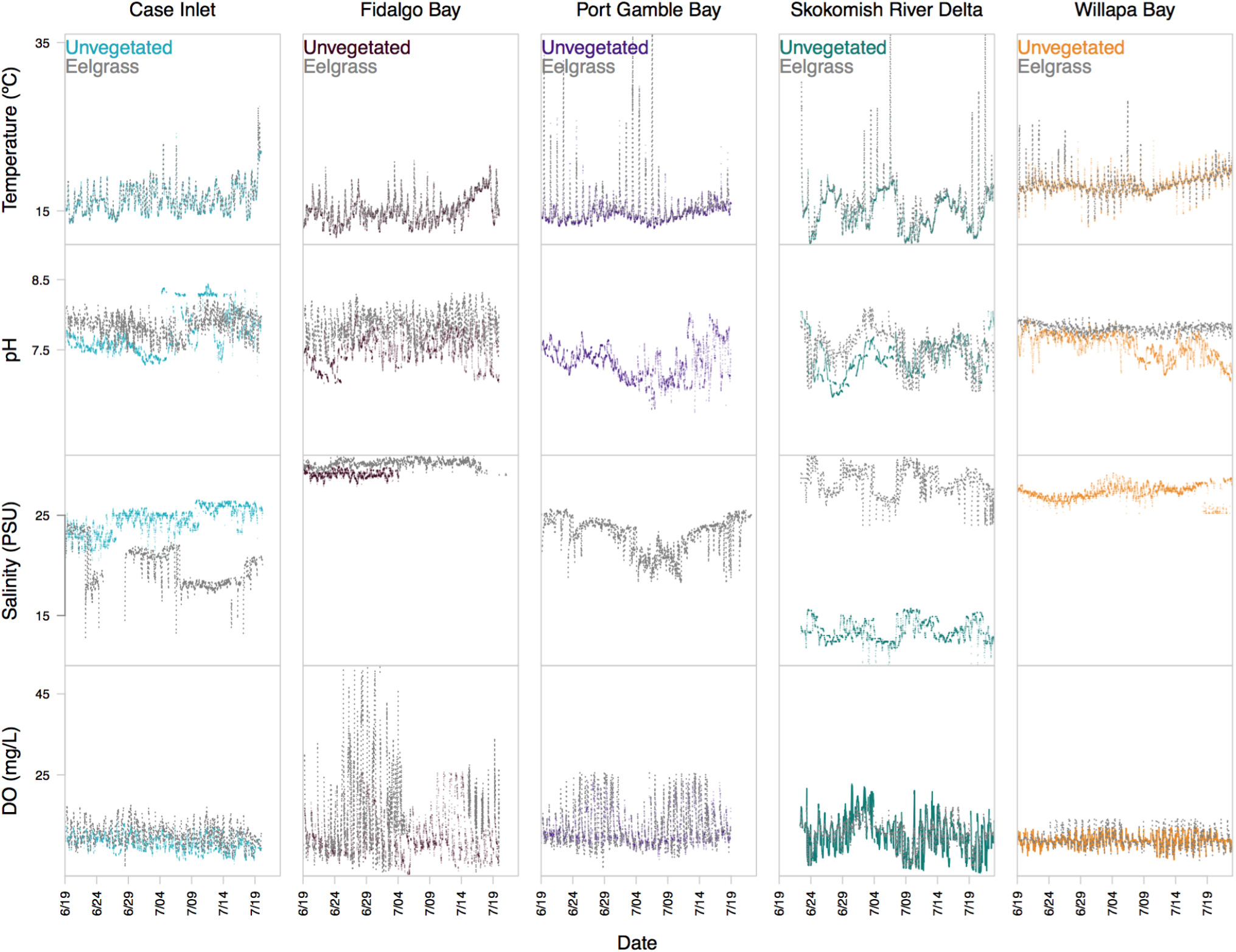
Environmental variables for each site (Case Inlet, Fidalgo Bay, Port Gamble Bay, Skokomish River Delta, and Willapa Bay) and habitat (unvegetated and eelgrass) over the course of the 29 day outplant.

**Table 3.** Environmental conditions (mean ± standard deviation) at outplant locations (Case Inlet (CI), Fidalgo Bay (FB), Port Gamble Bay (PG), Skokomish River Delta (SK), Willapa Bay (WB).

| Metric | CI | FB | PG | SK | WB |
| --- | --- | --- | --- | --- | --- |
| Mean Temperature (°C) | 16.1 ± 1.7 | 14.8 ± 1.8 | 15.1 ± 2.9 | 15.1 ± 2.2 | 18.0 ± 1.3 |
| Mean Salinity (PSU) | 24.47 ± 1.35 | 30.20 ± 0.38 | 23.25 ± 1.56 | 13.43 ± 1.04 | 27.28 ± 0.73 |
| Mean Dissolved Oxygen Content (mg/L) | 8.27 ± 1.91 | 9.63 ± 4.59 | 11.70 ± 3.70 | 9.99 ± 4.58 | 8.76 ± 1.78 |
| Mean pH | 7.63 ± 0.19 | 7.54 ± 0.23 | 7.33 ± 0.25 | 7.37 ± 0.24 | 7.56 ± 0.18 |

**Table 4.**
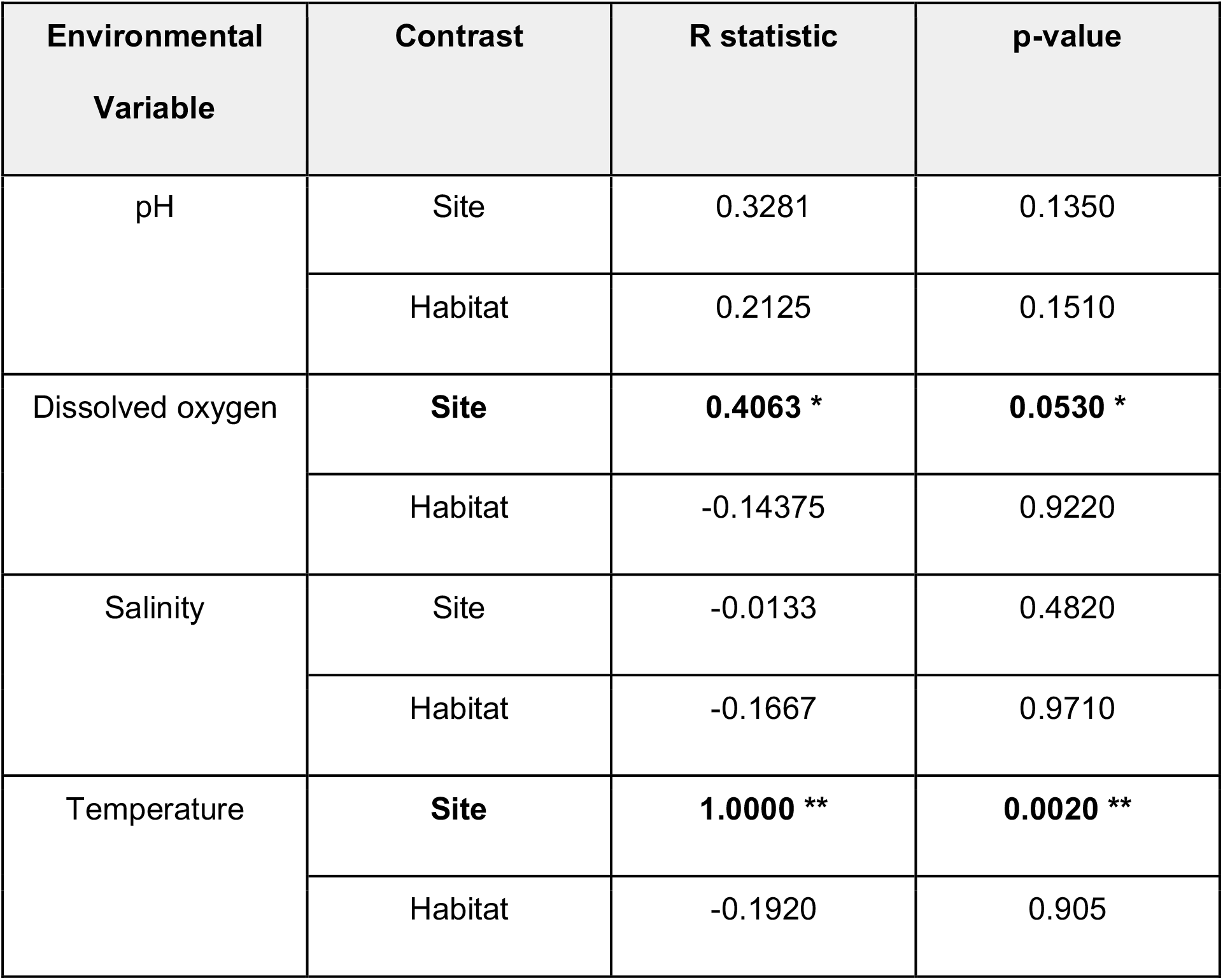
Analysis of Similarities (ANOSIM) R statistic and p-value for mean values of environmental variables by site and habitat. Values significant at the 0.05 level are marked by **, and values significant at the 0.10 level are marked by *.

| Environmental Variable | Contrast | R statistic | p-value |
| --- | --- | --- | --- |
| pH | Site | 0.3281 | 0.1350 |
|  | Habitat | 0.2125 | 0.1510 |
| Dissolved oxygen | **Site** | **0.4063 \*** | **0.0530 \*** |
|  | Habitat | -0.14375 | 0.9220 |
| Salinity | Site | -0.0133 | 0.4820 |
|  | Habitat | -0.1667 | 0.9710 |
| Temperature | **Site** | **1.0000 \*\*** | **0.0020 \*\*** |
|  | Habitat | -0.1920 | 0.905 |

**Table 5.** Analysis of Similarities (ANOSIM) R statistic and p-value for environmental variable variances by site and habitat. Values significant at the 0.05 level are marked by **, and values significant at the 0.10 level are marked by *.

| **Environmental Variable** | **Contrast** | **R statistic** | **p-value** |
| --- | --- | --- | --- |
| pH | **Site** | **0.5313 \*\*** | **0.0180 \*\*** |
|  | Habitat | -0.1813 | 0.9340 |
| Dissolved oxygen | **Site** | **0.6800 \*\*** | **0.0030 \*\*** |
|  | Habitat | -0.1240 | 0.8480 |
| Salinity | Site | **0.8400 \*\*** | **0.0130 \*\*** |
|  | Habitat | -0.1250 | 0.7080 |
| Temperature | Site | **0.9400 \*\*** | **0.0010 \*\*** |
|  | Habitat | -0.2200 | 0.9870 |

Although oyster position was standardized by tidal height, outplanted oysters experienced different amounts of exposure at low tide at each site due to differences in tidal amplitude (X4^2^ = 25.29, p < 0.0001). Oysters at FB spent the highest percent of time emersed at tides less than 1 foot MLLW (13.99%; 188 hours), followed by PG (10.93%; 146 hours), SK (10.29%; 138 hours), CI (9.35%; 125 hours), and WB (8.48%; 113 hours). CI and FB outplants (p = 0.0021), FB and SK outplants (p = 0.0324), and FB and WB outplants (p < 0.0001) spent significantly different amounts of time emersed during low tide conditions.

### Data Independent Acquisition Mass Spectrometry

Out of 39816 predicted proteins in the *C. gigas* FASTA proteome, 9047 proteins were detected in *C. gigas* across five sites and two habitats using DIA (Skyline auto peak picker error rate 24.3% ± 25%, range: 0% to 100%). Proteins detected included, but were not limited to, those annotated from processes such as responses to hypoxia and oxidative stress, removal of superoxide radicals, protein folding, muscle organ development, and negative regulation of apoptosis (Raw data: PeptideAtlas accession no. PASS01305).

### Selected Reaction Monitoring Assay

Differential abundance of protein targets was evaluated at the peptide level after combining technical replicates. Proteins were considered differentially abundant if at least one monitored peptide was differentially abundant. There was no significant difference in SRM peptide abundance between unvegetated and eelgrass habitats across sites (One-way ANOSIM; R = 0.0440, p = 0.1220). Abundance data from both habitats were pooled for downstream analyses comparing protein abundances among sites (Raw data: PeptideAtlas accession no. PASS01304).

Rank distances between peptide abundances in multivariate space revealed differences in peptide abundance between WB versus the other four sites (NMDS: stress = 0.0750, p = 0.0099). One-way ANOSIM by site demonstrated no significant differences in peptide abundance (R = 0.0640, p = 0.065), but a one-way ANOSIM by region (PS vs. WB) revealed a trend towards peptide abundance differences (R = 0.2260, p = 0.0530). Peptide abundance was significantly different between FB and Willapa Bay (Table 6; Pairwise ANOSIM; R = 0.2568, p = 0.0350). Environmental variables explained 29% of variance in peptide abundance, but the proportion of variation explained was not significant (ANOVA for RDA; F_6,19_ = 1.3064; p = 0.1950). Temperature mean and variance were the two most influential environmental predictors of peptide abundance, but were not significant at the 0.05 level (ANOVA on RDA; Table 7; temperature mean: F_1,19_ = 2.2375, p = 0.0650; temperature variance: F_1,19_ = 2.1627, p = 0.0670). Peptides differentially abundant between FB and WB are primarily positively loaded onto temperature mean, with two negatively loaded on temperature variance. Robustness of analysis was evaluated by performing a second RDA to predict peptide abundance based on mean and variance of temperature and pH at time of collection. Due to missing values, dissolved oxygen and salinity could not be included in analysis. Temperature variance was the most influential predictor of peptide abundance at time of collection, but was not significant at the 0.05 level (ANOVA on RDA; F_1,13_ = 2.2312, p = 0.0800).

**Table 6.** Results of pairwise ANOSIM tests between sites for protein abundance data. R values are listed above the diagonal, and p values are listed below the diagonal. Values significant at the 0.05 level are marked by **, and values significant at the 0.10 level are marked by *.

|  | CI | FB | PG | SK | FB |
| --- | --- | --- | --- | --- | --- |
| CI | - | 0.0197 | -0.0707 | -0.0583 | 0.0926 |
| FB | 0.3310 | - | -0.0313 | 0.0949 | 0.2568 |
| PG | 0.7800 | 0.5992 | - | -0.0061 | 0.0727 |
| SK | 0.7330 | 0.0860 | 0.3820 | - | 0.1547 |
| WB | 0.1600 | **0.0350 \*\*** | 0.192 | **0.0790 \*** | - |

**Table 7.** Significance of each predictor in the RDA. Salinity mean and variance were not included due to missing values. Predictors significant at the 0.10 level are marked by *.

| **Environmental Variable** | **Df** | **Variance** | **F** | **Pr(>F)** |
| --- | --- | --- | --- | --- |
| **Temperature Mean** | 1 | 0.0014 | **2.3275 \*** | **0.0650 \*** |
| **Temperature Variance** | 1 | 0.0013 | **2.1627 \*** | **0.0670 \*** |
| **pH Mean** | 1 | 0.0003 | 0.5264 | 0.7810 |
| **pH Variance** | 1 | 0.0007 | 1.2171 | 0.3010 |
| **Dissolved Oxygen Mean** | 1 | 0.0006 | 1.0354 | 0.3490 |
| **Dissolved Oxygen Variance** | 1 | 0.0003 | 0.5692 | 0.7250 |
| **Residual** | 19 |  |  |  |

Peroxiredoxin-5 (PRX), catalase (EC 1.11.1.6) (CAT), NAD(P) transhydrogenase (NADPt), glucose-6-phosphate 1-dehydrogenase (G6PD), carbonic anhydrase (CA), protein disulfide-isomerase 1 and 2 (PDI) had peptides that were identified as differentially abundant between Willapa Bay and Fidalgo Bay by SIMPER analysis (Figure 3). Fidalgo Bay peptide composition was differentiated by CA abundance, while all other significantly abundant proteins differentiated Willapa Bay. These proteins are involved in general cellular stress responses like reactive oxygen species neutralization or protein folding (Table 8). All differentially abundant proteins were detected at higher levels in the WB oysters than in oysters from the other four PS locations (CI, FB, PG, and SK), regardless of protein function (Figure 4). No differences in protein abundance were detected among the PS sites (Table 6).

**Figure 3.**
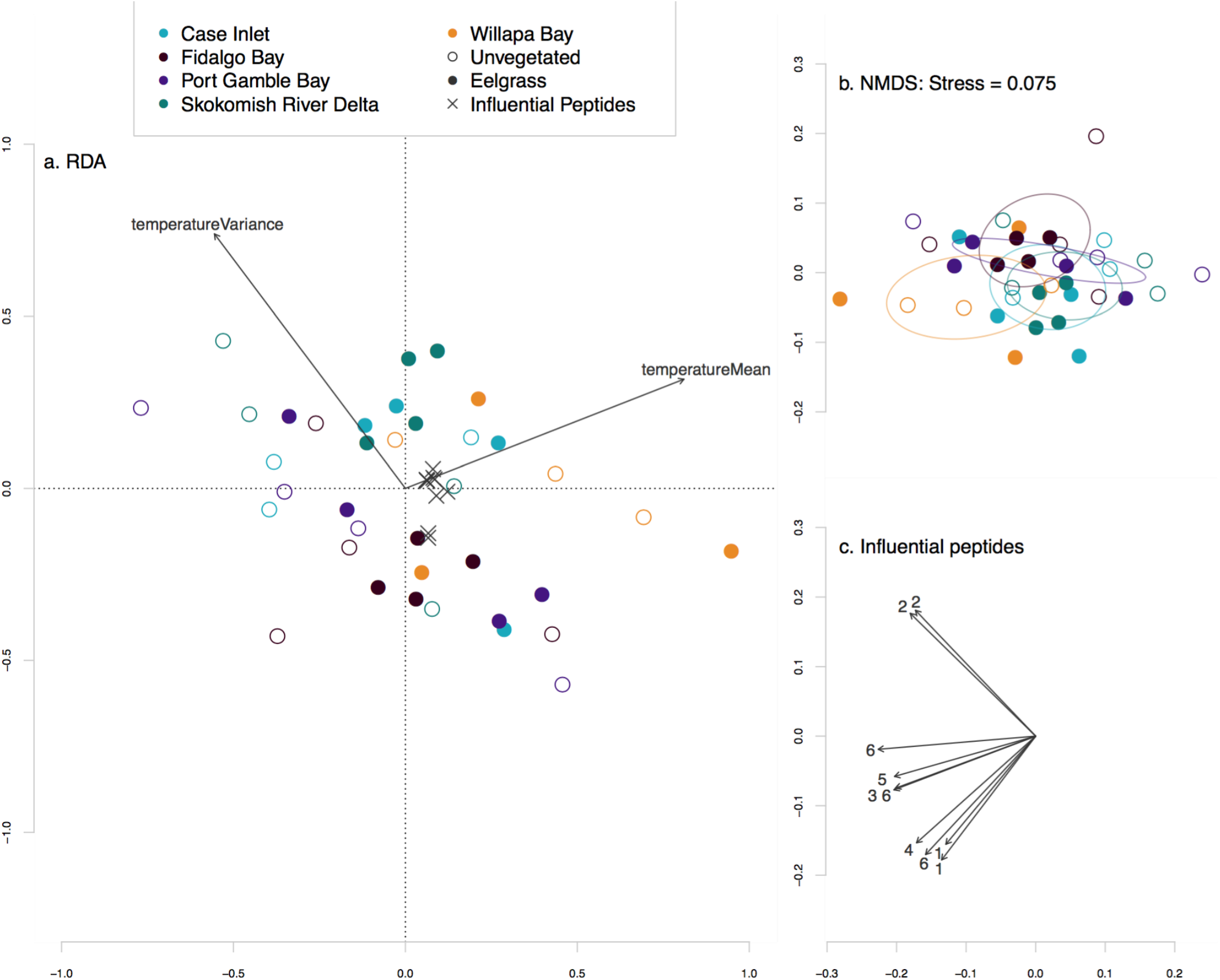
Ordination results for peptide abundance and environmental data. a) Environmental variables (pH, dissolved oxygen, salinity, and temperature) explained 30% of variance in peptide abundance data; however, redundancy analysis (RDA) for peptide abundance constrained by environmental variable was not significant (ANOVA; F_6,19_ = 1.306, p = 0.195). Temperature mean and variance were the most influential, yet nonsignificant, environmental predictors (ANOVA; temperature mean: F_1,25_ = 2.3275, p = 0.065; temperature variance: F_1,25_ = 2.1527, p = 0.067). Peptides differentially abundant between Fidalgo Bay (FB) and Willapa Bay (WB) are primarily positively loaded onto temperature mean, with two negatively loaded on temperature variance. b) Non-metric multidimensional scaling plot depicting peptide abundance by site and habitat, with 95% confidence ellipses around each site (stress = 0.075, p = 0.0099). c) Peptides that contributed to significant peptide abundance differences between FB and WB, as determined by post-hoc similarity percentage analysis (SIMPER). Peptides denoted 1 correspond to peroxiredoxin-5 (PRX), 2 for carbonic anhydrase (CA), 3 for catalase (CAT), 4 for glucose-6-phosphate 1-dehydrogenase (G6PD), 5 for NADP transhydrogenase (NADPt), and 6 for protein disulfide isomerases 1 and 2 (PDI). FB peptide composition was differentiated by CA abundance, while all other proteins influenced peptide abundances at WB.

**Figure 4.**
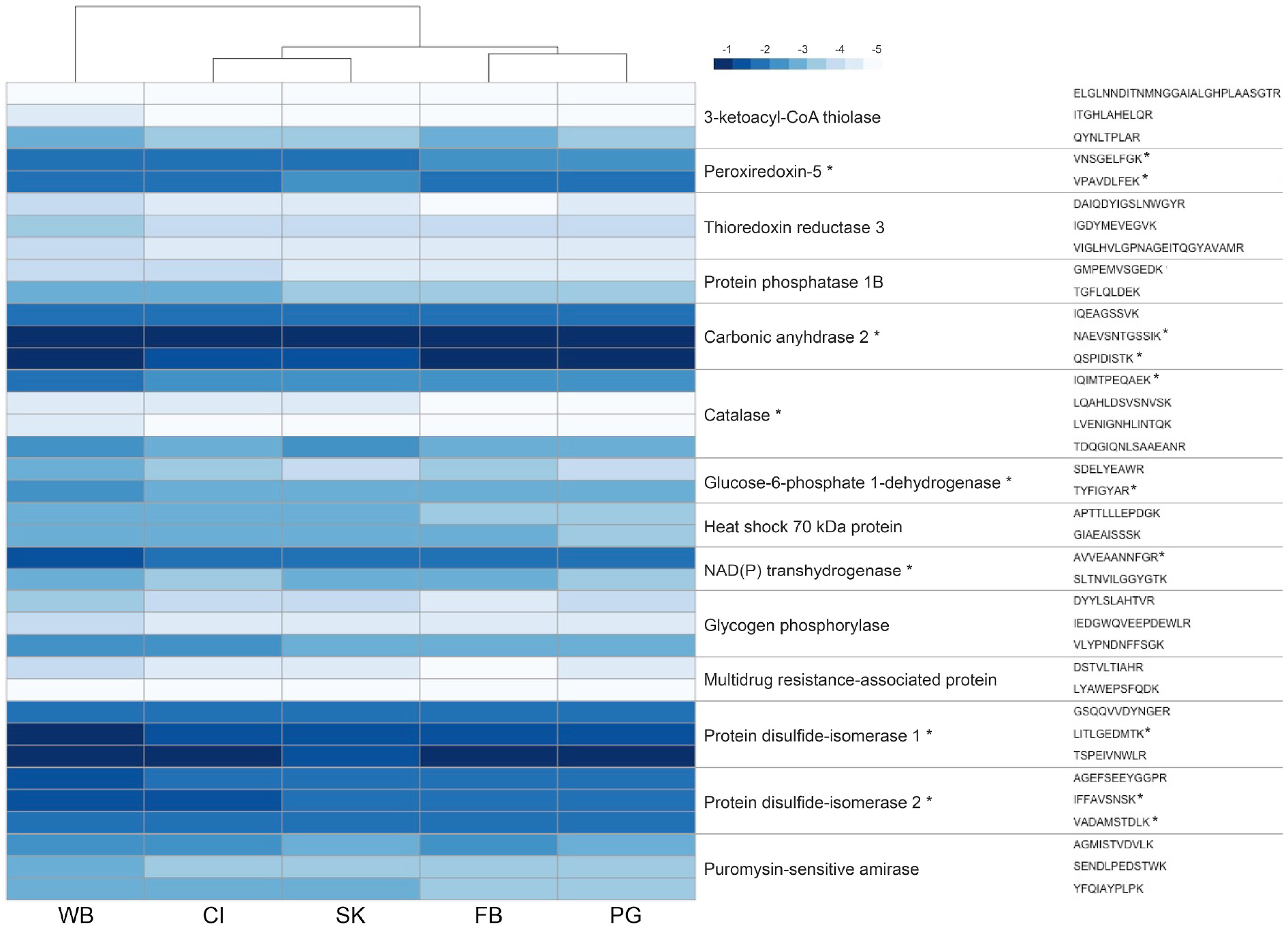
Average protein abundance by constituent peptides across experimental sites from SRM. Peptide abundance data at each site was averaged, then log transformed. Proteins were considered differentially abundant if at least one constituent peptide was significantly different (indicated by an asterisk). There were no significant differences in protein abundance among the Puget Sound locations, or between unvegetated and eelgrass habitats. Proteins were only significantly different between Fidalgo Bay and Willapa Bay or Skokomish River Delta and Willapa Bay.

**Table 8.** Functions of differentially abundant proteins

| **Protein** | **Function** | **Citations** |
| --- | --- | --- |
| Peroxiredoxin-5 (PRX) | Scavenges ROS located in mitochondria using | (Tomanek 2014;<br>Sussarellu et al. 2012) |
|  | cysteine residuals to<br>reduce substrates |  |
| Catalase (CAT) | Degrades hydrogen<br>peroxide into water and<br>oxygen | (Flores-Nunes, Mattos, et<br>al. 2015); (Sussarellu et al.<br>2012) |
| NAD(P) transhydrogenase<br>(NADPt) | Involved in maintenance of<br>cellular redox state | (Sussarellu et al. 2010,<br>2012) |
| Carbonic anhydrase (CA) | Maintenance of acid-base<br>balance by catalyzing CO<sub>2</sub><br>conversion to HCO<sub>3</sub><sup>-</sup> | (David et al. 2005; L. M.<br>Parker, Ross, and<br>O'Connor 2011; Wei et al.<br>2015) |
| Glucose-6-phosphate 1-dehydrogenase (G6PD) | Catalyzes first step in the<br>pentose phosphate<br>pathway and controls rate<br>of pathway's oxidative<br>portion | (L., n.d.) |
| Protein disulfide-isomerase<br>1 (PDI1) and Protein<br>disulfide-isomerase 2<br>(PDI2) | General chaperone protein<br>that aids in protein disulfide<br>bond formation and<br>isomerization | (Silvestre et al. 2006;<br>Vargas-Albores et al. 2009;<br>P. Zhang et al. 2014) |

## Discussion

Among sibling Pacific oysters placed at five sites in Washington State, we found higher abundances of seven proteins — antioxidant enzymes and molecular chaperones — in oysters outplanted at WB. Significant differences in protein abundances were detected between WB and either FB or SK; no differences were observed among PS sites. Increased antioxidant and molecular chaperone protein abundance are conserved stress responses consistently documented as responses to extreme temperatures, hyposalinity, low pH and air exposure (Tomanek 2014; Zhang et al. 2015). Warmer and less variable temperature conditions at WB may have driven differential protein abundances (Figure 3). Validation of RDA results by correlating peptide abundance to environmental conditions at the time of collection demonstrates that proteomic methods are suitable for evaluating the influence of month-long environmental variation on organismal physiology.

To our knowledge, this is the first time combined use of DIA and SRM analytical methods have been applied in an ecological context. Previously, both gel-based (Parker et al. 2011; Zhang et al. 2015; Campos et al. 2016) and gel-free (Dineshram et al. 2016; Timmins-Schiffman et al. 2017; Müller et al. 2018) proteomic methods have been used in lab settings to evaluate the effects of environmental conditions on organismal physiology, with gel-based methods having a limited dynamic range. Similar gel-free methods have also been used to examine physiology of other non-model marine species (Plumel et al. 2013; Kültz et al. 2015). Our methods allowed for increased acquisition of proteomic data for a non-model organism. We detected over 9000 proteins with DIA — far more than previous studies have detected. The application of a novel, two step, gel-free proteomic approach broadened the scope of proteins detected in the DIA phase, thus revealing more possibilities for SRM target design and also completing the first step for future proteomic studies in this species. The SRM assay allowed investigation of specific biomarkers with a high-throughput analysis.

### Antioxidant enzymes and Acid-Base Regulation

Higher antioxidant enzyme abundances can be a direct response to an increase in reactive oxygen species (ROS) (Limón-Pacheco & Gonsebatt 2009; Zhang et al. 2015). During electron transport, oysters can produce ROS that induce oxidative stress if not neutralized (Abele et al. 2007; Limón-Pacheco & Gonsebatt 2009). Peroxiredoxin-5 (PRX), and catalase (CAT) scavenge these ROS and degrade them before they cause cellular harm, while NAD(P) transhydrogenase (NADPt) maintains the cellular redox state (Limón-Pacheco & Gonsebatt 2009; Sussarellu et al. 2012; Flores-Nunes et al. 2015). Higher abundances of antioxidant enzymes in WB oysters suggests the need for ROS scavenging to acclimatize to local conditions.

Elevated antioxidant enzyme abundance at WB may be a response to higher levels of ROS brought on by heat stress. Mollusc species, like *C. gigas*, have been known to increase ROS production at higher temperatures (Abele et al. 2007; Tomanek 2014). Warmer and more variable temperature conditions at WB could be driving ROS production and in turn, higher abundances of PRX, CAT, and NADPt to ameliorate ROS-associated stress (Zhang et al. 2015). The shallow bathymetry of WB may have also contributed to elevated ROS presence and the need for antioxidant enzymes. At low tide, shallow waters would warm faster, leading to the higher temperatures observed at WB (Table 1). Warmer waters at low tide may prompt oysters to spend more time with their shells closed, inducing hypoxia and hypercapnia. Low oxygen concentrations within the shell could then augment ROS levels (Guzy & Schumaker 2006). Oysters could respond through increased abundance of antioxidant enzymes (Sussarellu et al. 2012).

The need for WB outplants to regulate internal acid-base conditions is demonstrated by higher abundance of carbonic anhydrase (CA) in these oysters. *C. gigas* can regulate hemolymph pH by increasing HCO_3_^−^ concentration via the conversion of CO2 to HCO_3_^−^, catalyzed by CA (Michaelidis et al. 2005; Wei et al. 2015). If warmer water conditions at WB prompted shell closure, *C. gigas* would need to switch to anaerobic metabolism, inducing a significant reduction in hemolymph pH (Michaelidis et al. 2005). Buffering of hemolymph pH could be accomplished by elevated CA abundance. Although oysters at FB spent more time in low tide conditions that would also prompt shell closure and similar molecular responses, upregulated protein abundance at WB implies conditions at this site required a greater proteomic response in these specific biomarkers for acclimatization.

Higher abundance of glucose-6-phosphate 1-dehydrogenase (G6PD) in WB oysters indicates these oysters maintained metabolic activity in warmer temperature conditions. G6PD catalyzes the oxidative portion of the pentose phosphate pathway, and products from this pathway are often the precursors for nucleic and aromatic amino acids (Livingstone 1981). Additionally, G6PD activity generates NADPH and can indirectly regulate the redox environment and ameliorate effects of ROS (Livingstone 1981). For example, exposure to ROS-inducing pollutants led to increased G6PD activity in *C. gigas* ctenidia (Flores-Nunes et al. 2015). Increased abundance of G6PD in WB not only could have maintained transcription and translation processes, but also levels of cellular and metabolic activity by indirectly dealing with ROS.

ROS are produced in response to many environmental changes, thus biomarkers of ROS scavenging are difficult to link to a single environmental difference in a variable and complex setting. For example, reduction of ROS was found to be a common response to both increased temperatures and aerial exposure in *C. gigas* (Zhang et al. 2015). Since ROS mediation is a conserved response to several stressors, it is possible that environmental parameters we did not measure (e.g., contaminants, microbiota abundance, trace metals), or a combination of environmental parameters, could explain the observed variation in antioxidant enzyme abundance. Future work at these locations should take these variables into account.

### Molecular Chaperones Involved in Protein Folding

Much like cellular response to ROS, increased levels of protein disulfide isomerase 1 and 2 (PDI) demonstrate a generalist response to conditions at WB. PDI is a general chaperone protein that forms disulfide bonds and assists with protein folding (Vargas-Albores et al. 2009). Higher PDI abundance would reflect a need to refold misshapen proteins. In this regard, PDI would function similarly to molecular chaperones like heat shock proteins. Several invertebrate taxa demonstrate higher PDI abundance when faced with an immune challenge or metal contamination. When faced with an immune challenge, Pacific white shrimp (*Litopenaeus vannamei*) hemocytes synthesized higher abundances of immune response proteins, followed by elevated abundance of PDI to correct disulfide bonds in these proteins (Vargas-Albores et al. 2009). An immune challenge could also lead to more misfolded proteins that PDI would need to refold to avoid cellular damage (Zhang et al. 2014). In flat oysters (*Ostrea edulis*), PDI expression increased in response to disseminated neoplasia (Silvestre et al. 2006; Martín-Gómez et al. 2013). Metal contamination at WB could also elevate PDI abundance: Chinese mitten crabs (*Eriocheir sinensis*) chronically exposed to cadmium over-expressed PDI (Silvestre et al. 2006; Martín-Gómez et al. 2013). Since neither disease burden or environmental contamination was measured in this study, it is impossible to know if either triggered the PDI response. Examination of these factors in future studies may elucidate the specifics of elevated PDI abundance in WB.

### Proteomic Responses in Puget Sound and Between Habitats

Due to the observed differences across environmental parameters between Willapa Bay and Puget Sound locations, similar abundances for proteins involved in various environmental responses may be evidence of physiological plasticity. One particular protein that we expected to be differentially abundant was heat shock 70 kDA protein 12A (HSP12A), since Willapa Bay had higher average temperatures (Table 3). However, this trend was not observed. Higher abundances of heat shock proteins (HSPs) are generally induced when organisms are exposed to thermal stress (Hamdoun et al. 2003; Zhang et al. 2015). In our experiment, average temperatures *C. gigas* experienced were lower than the 30°C threshold to induce elevated HSP expression in a controlled setting (Figure 2; Table 3), which would explain why we did not see elevated HSP12A expression in WB (Hamdoun et al. 2003).

The lack of differential abundance for protein targets — both among Puget Sound sites and between unvegetated and eelgrass habitats — was unexpected. These similar proteomic profiles could be due to four factors. First, it is possible that a different suite of environmental response proteins in the SRM assay could have yielded a different view of acclimatization to the various outplant sites; however, the targets we chose have proven to yield insight into a range of environmental responses in previous studies (ex. Table 8). Second, outplant duration could have been too short to capture varied physiological response within Puget Sound, or oysters could have also acclimatized to conditions in their outplant locations. Third, post-translational modifications may have influenced how we detected proteins. Abundance between sites or habitats may have been similar, but addition of post-translational modifications may have varied. Finally, it is possible that the proteomic response was not different because the environmental conditions that would elicit up- or down-regulation of monitored proteins were similar across these five locations. For example, we found no differences in environmental conditions between eelgrass and unvegetated areas, nor any differences in protein responses between these habitats (Table 4; Table 5). Our results suggest that the potential vegetation-based systems of remediation could be limited under current conditions in the field. A recent comparison of stable isotopes and fatty acid composition in *C. gigas* at eelgrass and unvegetated habitats found no differences in δ^13^C, δ^15^N, total fatty acids, or proportional fatty composition in WB outplants, providing another line of evidence that suggests habitat may not affect *C. gigas* physiological performance (Lowe et al. 2018).

## Conclusion

Differential abundance of target proteins observed between sibling oysters placed for 29 days in WB or PS indicates that environmental factors at WB tended to increase antioxidant enzyme and molecular chaperone protein abundance in Pacific oysters. This study is one the first to link regional environmental conditions to physiological responses in Puget Sound and Willapa Bay, as well as compare responses between Puget Sound and Willapa Bay. Understanding the difference between these two estuaries is important for the persistence of oyster reefs and aquaculture in the face of climate change. Of the environmental parameters measured, higher mean temperature, as well as less variable temperature conditions, at Willapa Bay may have influenced protein abundance. The lack of protein abundance differences between PS sites may imply that 2016 conditions were well-within the tolerances of outplanted *C. gigas*, so patterns of stress response witnessed under laboratory conditions were not apparent in the field. Together, the results generate a suite of new hypotheses for lab and field experiments comparing the effects of environmental conditions on physiological responses of marine invertebrates.

As global climate change continues to rapidly influence estuarine dynamics, this study illustrates the importance of investigating the effect of multiple modes of change on organismal physiology *in situ*. Our finding that oysters used generalist proteins to ameliorate stressors implies that assays for elucidating responses to multiple drivers *in situ* should include these conserved responses in addition to specific, stressor-related proteins identified in laboratory experiments. Pacific oysters are grown commercially throughout the Pacific Northwest region of the United States, so it is possible the population used for this study has a broad environmental tolerance. Proteomic assays allow for quantification of that tolerance, which is crucial for aquaculture industry and natural resource management. Future *in situ* explorations of environmental drivers on physiology should include a longer outplant duration, additional environmental variables, and multiple sampling points to provide a clearer connection between ecosystem dynamics and physiological performance.

## Acknowledgements

This work was funded by the Washington State Department of Natural Resources Interagency Agreement 16-241 and Washington Sea Grant (Award: NA14OAR4170078; Project R/SFA-8). Brittany Taylor (Taylor Shellfish) provided *C. gigas* for the outplant experiment. The University of Washington Proteomic Resource and Priska von Haller assisted with mass spectrometry. Jarrett Egertson (Department of Genome Sciences) helped design the DIA acquisition. Austin Keller (Department of Genome Sciences) adapted MSConvert to correctly demultiplex and convert DIA data files. Sean Bennet (School of Aquatic and Fishery Sciences), and Han-Yin Yang and Brian Searle (Department of Genome Sciences) assisted with running Pecan. We also thank three anonymous reviewers for their valuable feedback on the manuscript.

## Supplementary Material

Raw data can be accessed in the PeptideAtlas under accession PASS01304 and PASS01305. Skyline documents can be found on Panorama Public.

## Abbreviations

PS: Puget Sound
CI: Case Inlet
FB: Fidalgo Bay
PG: Port Gamble Bay
SK: Skokomish River Delta
WB: Willapa Bay
DIA: Data Independent Acquisition Mass Spectrometry
SRM: Selected Reaction Monitoring Assay

